# Decoding natural scenes from patterned optogenetic responses in mouse visual cortex

**DOI:** 10.64898/2026.09.21.753135

**Authors:** Shanshan Jia, Tianyu Xin, Chunpeng Li, Haofu Ji, Ruixiang Wu, Shu Wang, Qingchun Guo, Zhaofei Yu, Jian K. Liu, Ya-tang Li

## Abstract

A central challenge in developing visual cortical prostheses is to determine how visual stimuli should be transformed into effective patterns of cortical stimulation. Although advances in stimulation technologies, including optogenetics, provide increasingly precise control over cortical activity, it remains unclear whether artificially evoked activity can reproduce the information content of naturally evoked visual representations. Here we establish a quantitative framework for evaluating visual encoding strategies by decoding cortical responses evoked by natural vision and patterned optogenetic stimulation. We developed a novel dual-modal paradigm in awake mice to bridge the gap between endogenous photostimulation and artificial network driving. By co-expressing the high-performance calcium indicator GCaMP6s and the red-shifted, ultra-sensitive opsin rsChRmine-oScarlet in the primary visual cortex (V1), we successfully translated dynamic natural movie frames into patterned, spatiotemporal optogenetic stimulation. Quantitative comparisons of macro-scale dynamics demonstrated that this patterned optogenetic injection evokes cortical states highly comparable and representationally aligned with those driven by actual visual photostimulation. To systematically evaluate the fidelity of these responses, we developed STAR, a deep learning model featuring spatial and temporal attention mechanisms, and successfully reconstructed the frames of natural movies from V1 signals under both experimental modalities. Collectively, our results demonstrate that complex sensory information can be both naturally encoded and synthetically injected into V1 circuits with high decoding fidelity. This work provides an empirical and computational proof-of-concept for intelligent, closed-loop biomimetic encoders, establishing a robust framework for next-generation cortical visual neuroprostheses and bidirectional brain-machine interfaces.

## 1 Introduction

While normal visual perception is driven by light entering the eye, visual cortical prostheses (VCPs) offer a promising solution for restoring vision in blind individuals by directly stimulating the visual cortex. The concept of cortical visual stimulation was proposed by Benjamin Franklin more than two centuries ago, and despite decades of experimental and clinical research, progress toward practical visual restoration remains limited (1; 2; 3; 4; 5; 6). Current efforts to advance the field have been pursued from two complementary directions. On one hand, substantial progress has been made in stimulation technologies, including electrical interfaces with increasingly high spatial resolution (7) and, more recently, optogenetic stimulation approaches that offer cell-type specificity and finer spatial control (8; 9). On the other hand, the algorithms that transform visual input into effective cortical stimulation patterns remain a major bottleneck for achieving meaningful artificial vision (10).

A fundamental challenge is determining whether an encoding algorithm preserves the visual information required for perception. In clinical practice, the effectiveness of VCPs is assessed primarily through subjective perceptual reports from implanted blind individuals. One alternative strategy is to treat neural activity as an information-bearing representation and assess its fidelity by decoding the original visual input. If visually evoked and stimulation-evoked population responses support comparable decoding performance, they are likely to contain similar visual information. Such a decoding-based framework provides a quantitative and task-relevant metric for evaluating visual encoding strategies, as demonstrated in neuroprosthetics strategies of retinal vision restoration (11; 12; 13; 14; 15; 16).

Developing this framework requires measuring cortical responses to naturalistic visual inputs while reproducing comparable activity through artificial stimulation. While traditional studies have long relied on simplified, low-dimensional visual stimuli like moving gratings or localized spots of light, natural scenes and dynamic movies provide a more natural, relevant window into sensory processing of visual encoding and decoding (17; 18; 19; 20; 21; 22; 23). Meanwhile, recent advancements in mesoscopic wide-field calcium imaging now enable large-scale monitoring of neocortical dynamics, offering an unprecedented view of how distributed neural networks encode complex, non-stationary visual information across multiple interconnected cortical areas simultaneously (24). Reproducing these high-dimensional activity patterns with artificial stimulation, however, remains technically challenging. Conventional optogenetic stimulation typically employs spatially uniform or rudimentary patterned stimulation, which fails to capture the structured spatiotemporal dynamics of natural visual responses (25).

To bridge this critical gap, we establish a quantitative framework for evaluating visual encoding algorithms by comparing visual-evoked and optogenetic-evoked cortical responses to natural movies. To enable simultaneous patterned optogenetic stimulation and large-scale calcium imaging, we developed a dual-modal optical system utilizing the co-expression of the green calcium indicator GCaMP6s and the red-shifted, ultra-sensitive opsin (26; 27; 28). In this study, we systematically evaluated whether these synthetic, optogenetically driven cortical responses can effectively mimic the endogenous neural representations induced by actual visual photostimulation in awake mice. To evaluate the fidelity, constraint boundaries, and information content of these evoked cortical responses, we developed a deep learning framework, termed *STAR*, a deep learning model utilizing *S*patial and *T*emporal *A*ttention mechanisms for visual scene *R*econstruction, to decode and reconstruct the original natural movies directly from V1 wide-field calcium signals. We show that STAR successfully reconstructs frames of complex natural movies from both visual-evoked and optogenetic-evoked cortical responses. By demonstrating that complex sensory information can be both naturally encoded and artificially introduced into cortical networks with high decoding fidelity, our work provides a proof-of-concept framework that integrates neural decoding with patterned cortical stimulation and opens new avenues for studying mesoscale cortical computation.

## 2 Results

### 2.1 A dual-modal optical system for recording visually and optogenetically evoked responses to natural movies in mouse V1

To directly compare visually and optogenetically evoked responses from the same cortical neurons, we developed a dual-modal optical system for simultaneous wide-field calcium imaging and patterned optogenetic stimulation (Figure 1A). By virally expressing the red-shifted opsin rsChRmine in GP4.3 mice, which genetically express the calcium indicator GCaMP6s (Figures 1B–C), we directly measured both visual and optogenetic responses from the same neuronal population. For each animal, we first mapped retinotopic organization to locate V1 (Figures 1D–E), then presented a library of 20 natural movies spanning diverse visual scenes (Table S1) to evoke neural responses across V1.

**FIGURE 1.**
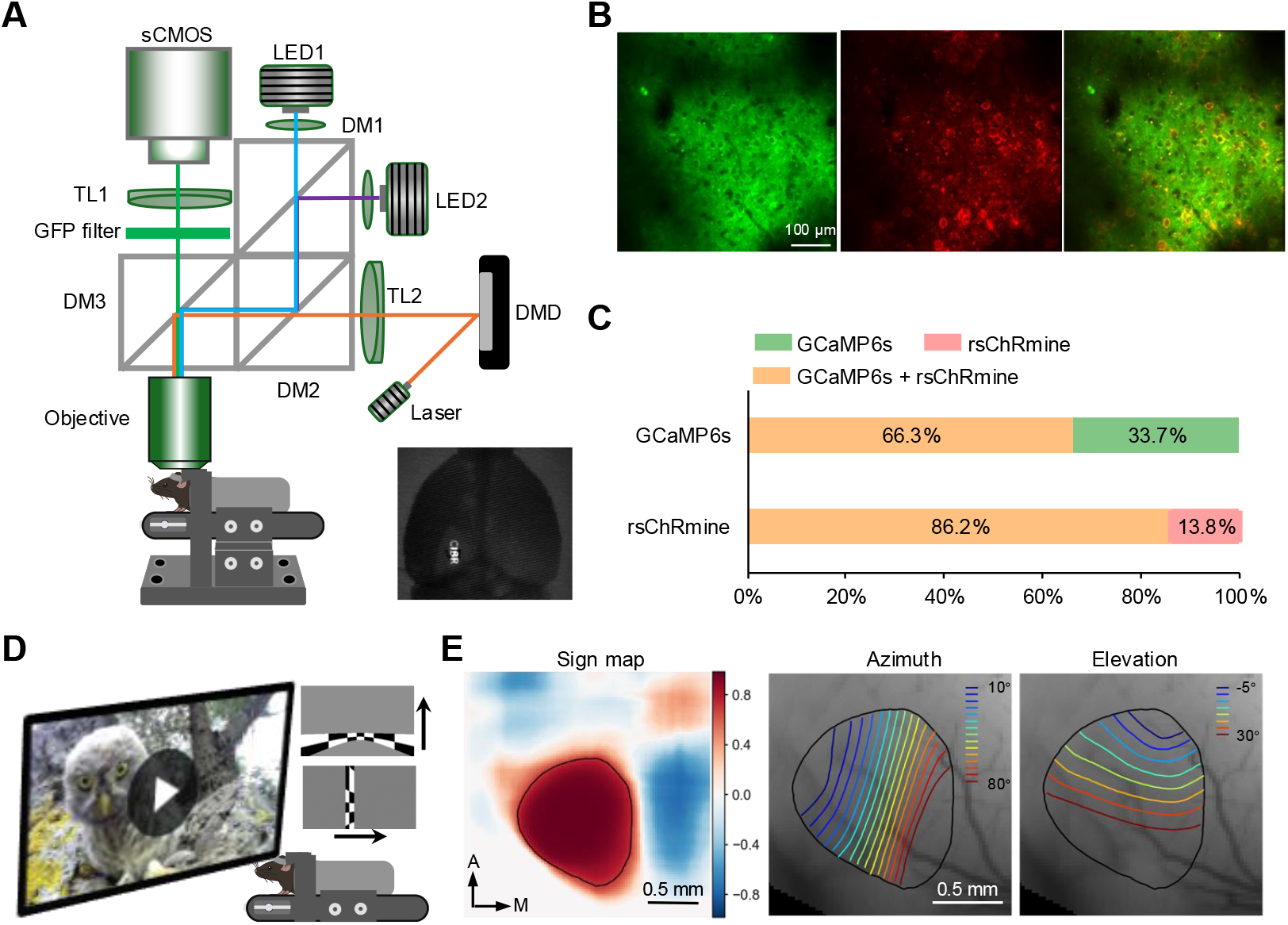
Experimental setup. (A) Schematic of the dual-modal system enabling simultaneous wide-field calcium imaging and patterned optogenetic stimulation. Inset, example patterned stimulation pattern. (B) Co-expression of GCaMP6s (green) and rsChRmine-oScarlet (red) in the mouse visual cortex. (C) Percentage of double-labeled neurons among GCaMP6s-positive neurons (*n*=169) and rsChRmine-positive neurons (*n*=130). (D) Visual stimulation paradigm. Visual stimuli include natural movies and drifting bars used for retinotopic mapping. (E) An example field sign map and corresponding retinotopic maps of azimuth and elevation.

To generate corresponding optogenetic stimulation patterns, we characterized cortical responses across a range of laser intensities and established the relationship between stimulation power and neural activity (Figures 2A). We next evaluated three feature-encoding strategies based on motion, object boundaries, and visual saliency. For each strategy, we quantified the similarity between visually and optogenetically evoked population responses using cosine similarity, Pearson’s correlation coefficient, and the soft F1 score. Among the three approaches, visual saliency consistently yielded the highest similarity and was therefore selected for all subsequent experiments (Figure 2B). Finally, we recorded visual-evoked and optogenetic-evoked population responses to natural movies for subsequent decoding analyses (Figures 2C and S1).

**FIGURE 2.**
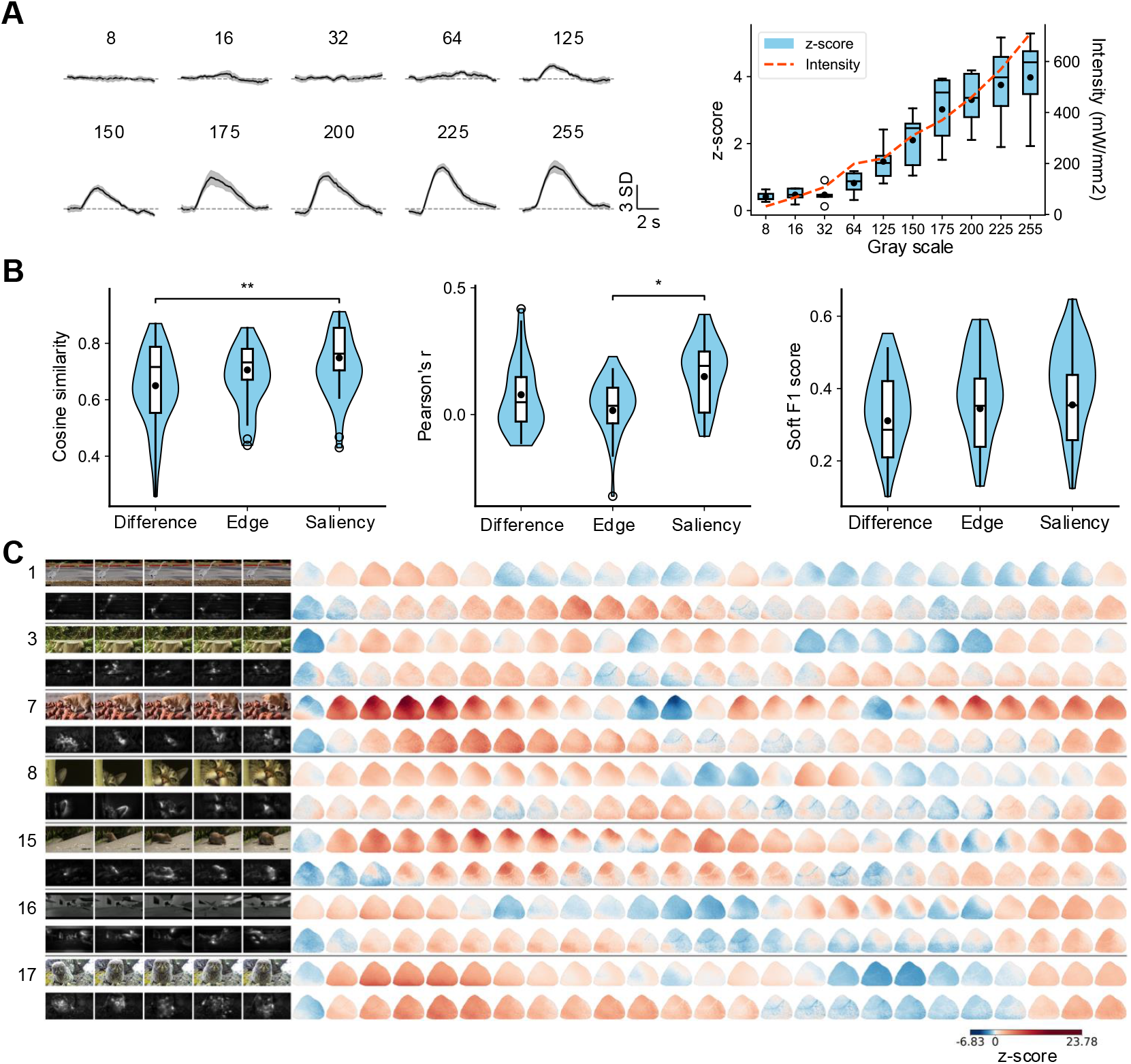
Performance comparison of encoding strategies. (A) Left, example z-scored calcium traces at different greyscale values. Right, relationship between grayscale value (LED intensity) and optogenetic-evoked calcium responses. *n* = 5 mice. (B) Comparison of encoding strategies based on the similarity between visually and optogenetically evoked responses, quantified using cosine similarity, the soft F1 score, and Pearson’s correlation coefficient. *n* = 40 recordings (20 videos from 2 mice). (C) Example visually and optogenetically evoked neural responses to natural movies in awake mouse V1. Left, clips of nature movies (odd rows) and corresponding optogenetic stimulation patterns (even rows). Right, evoked neural responses in V1 measured using wide-field calcium imaging.

### 2.2 STAR reconstructs natural visual scenes from visual-evoked and optogenetic-evoked V1 activity

To examine whether natural visual scenes viewed by awake mice can be decoded from V1 wide-field calcium dynamics, we developed STAR to reconstruct pixel-level visual stimuli (Figure 3A). The model employs a dual-branch architecture comprising modality-specific encoders paired with a shared reconstruction decoder. Visual-evoked and optogenetic-evoked signals are routed through dedicated encoder branches to capture and preserve their distinct response distributions. The extracted features are subsequently mapped into a unified latent representation space, from which the shared decoder reconstructs the visual images. This dual-encoder strategy effectively accommodates domain-specific variance across stimulation types, while the shared decoder forces the network to isolate common representational manifolds linked directly to visual content. Crucially, during inference, the model operates flexibly on unimodal V1 inputs; paired multi-modal signals are not required to generate accurate reconstructions.

**FIGURE 3.**
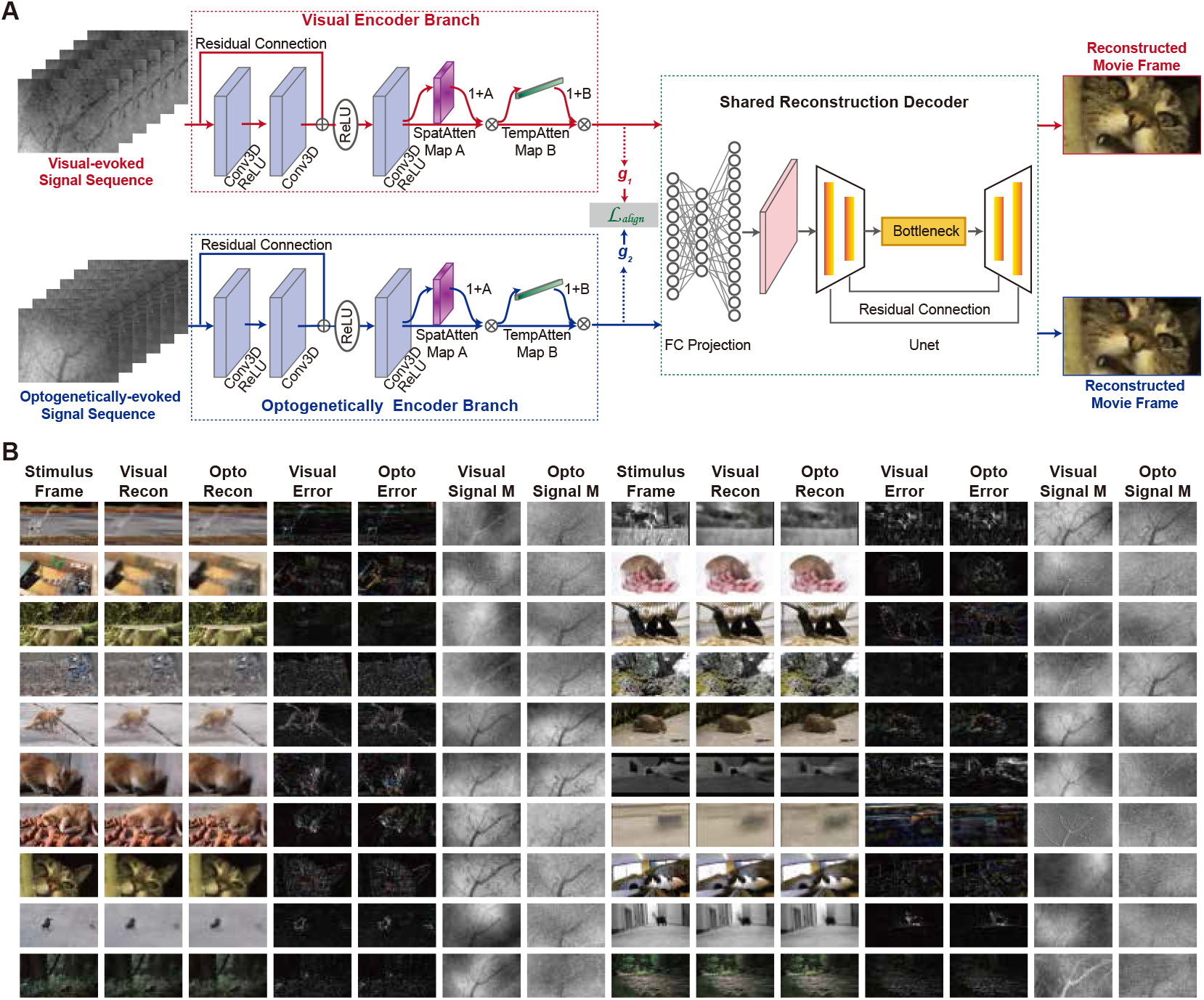
STAR for visual reconstruction. (A) Modality-specific encoders extract visual and optogenetically evoked V1 features, which are aligned and processed by a shared reconstruction decoder. Either modality can independently support reconstruction during testing. (B) Reconstructions, error maps, and V1 signal maps for 20 natural visual stimuli. Both modalities produce consistent visual content despite their different response distributions.

STAR achieved exceptional reconstruction performance across 20 dynamic natural visual stimuli (Figure 3B). Reconstructions generated independently from visual-evoked and optogenetic-evoked V1 inputs demonstrated striking qualitative consistency. Despite clear spatial differences in the raw V1 activity maps across modalities, both input streams yielded consistent reconstructions regarding object locations, overall scene layout, and major luminance structures. For scenes containing animal subjects, natural backgrounds, or distinct structural contours, both visual and optogenetical inputs successfully recovered the primary visual features of the original stimuli. These findings indicate that STAR successfully reconciles modality-specific discrepancies by mapping disparate neural dynamics into a common visual feature space.

Compared with existing baseline architectures, STAR demonstrated superior qualitative reconstruction fidelity, which was quantitatively confirmed by higher pixel-level fidelity (PSNR) and structural similarity (SSIM; Figure S2), even with single-trial recordings (Table S2). Ablation experiments confirmed that both spatial and temporal attention modules independently contribute to these performance gains (Table S3). Furthermore, training STAR on signals extracted from non-V1 cortical regions resulted in severe reconstruction degradation (Figure S3), proving that the decoding pipeline relies specifically on retinotopically localized visual responses rather than non-specific global cortical dynamics. Together, these results establish that STAR can efficiently and effectively decode complex natural visual scenes from mesoscopic V1 calcium signals in awake mice.

### 2.3 Stimulus content and visual category dependence of reconstruction fidelity

We further examined how reconstruction accuracy varied across diverse stimulus contents. Figure 4A shows the distributions of SSIM and PSNR across 20 natural video stimuli. Reconstruction performance differed substantially across movies. Certain stimuli consistently yielded higher SSIM and PSNR values, indicating that their primary spatial structures were more reliably recovered. In contrast, other stimuli exhibited lower performance scores and greater variance, indicating that STAR reconstruction fidelity is significantly modulated by specific visual content. Qualitative inspection reveals that stimuli characterized by prominent object structures or stable spatial layouts were reconstructed with higher fidelity. For example, video clips featuring distinct animal bodies or uncluttered backgrounds effectively preserved object localization and primary contours in the reconstructed frames. Conversely, scenes with complex backgrounds, indistinct boundaries, small targets, or rapid local movement presented greater reconstruction challenges, resulting in lower SSIM and PSNR metrics. These findings suggest that the amount of visual information stably encoded within V1 population activity directly constrains final reconstruction quality.

**FIGURE 4.**
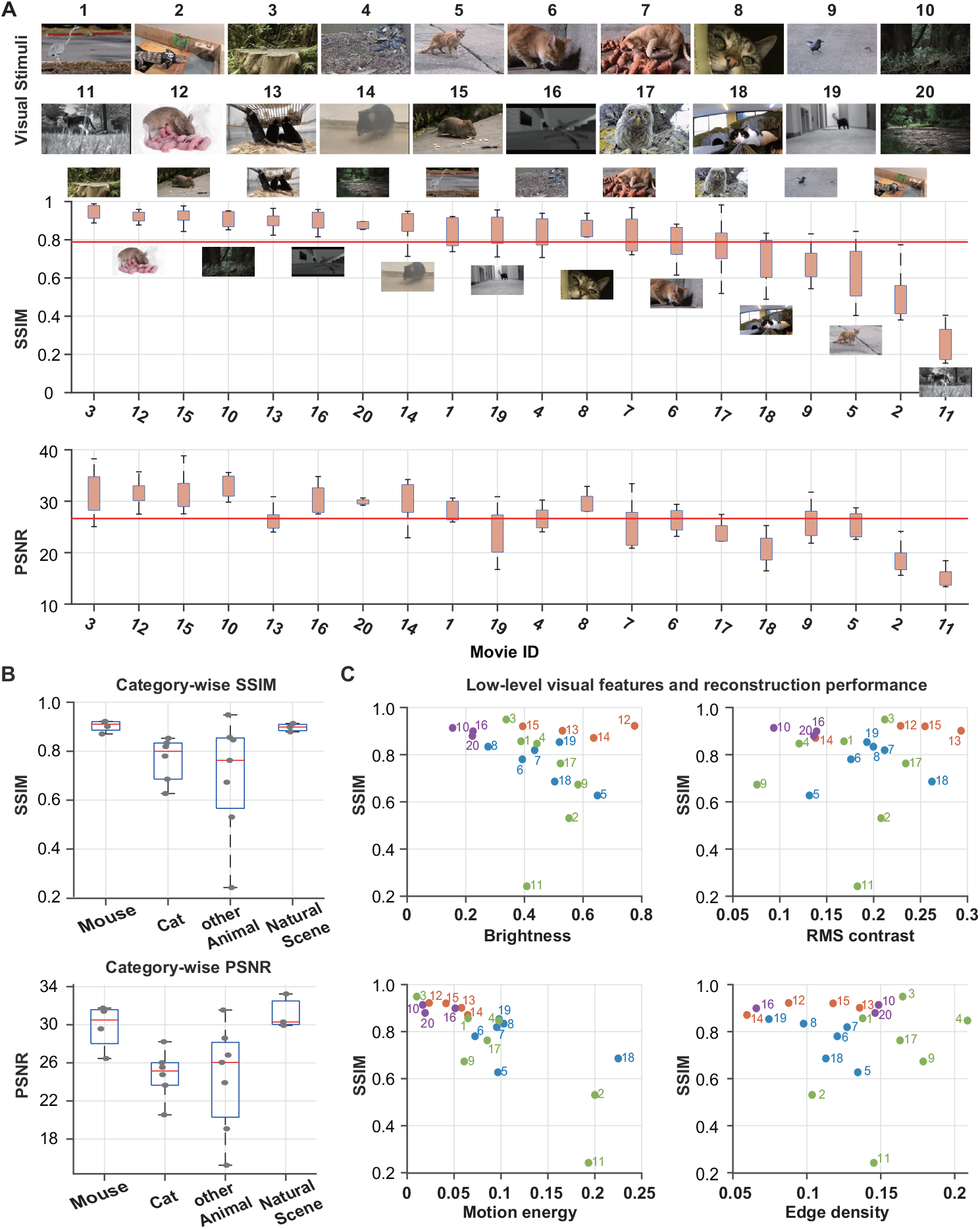
Reconstruction accuracy varies across stimulus content and visual categories. (A) SSIM and PSNR distributions across 20 natural video stimuli, sorted by reconstruction performance. Representative frames are shown above each movie ID. (B) Category-wise comparison of reconstruction performance for mouse, cat, other animal, and natural scene stimuli. (C) Relationship between reconstruction accuracy and low-level visual features, including brightness, RMS contrast, motion energy, and edge density. Each point represents one movie.

We next evaluated reconstruction performance across distinct visual categories. As shown in Figure 4B, SSIM and PSNR distributions varied across semantic groups. The mouse and natural scene categories displayed higher, more compact performance distributions, reflecting consistent structural recovery. Conversely, the cat and other animal categories exhibited broader variability, particularly within the other animal group, where reconstruction accuracy varied markedly across individual video samples. These indicate that decoding accuracy depends not only on high-level categorical labels, but also on the unique structural properties of each specific stimulus.

To isolate the underlying drivers of this variability, we analyzed the relationship between low-level visual features and reconstruction performance. Figure 4C compares SSIM against luminance, root-mean-square (RMS) contrast, motion energy, and edge density. The analysis demonstrates that no single low-level feature fully accounts for the variance in reconstruction accuracy. Stimuli featuring high motion energy or dense edge distributions frequently displayed variable reconstruction scores, suggesting that while low-level complexity contributes to decoding difficulty, it does not act as the sole determinant of performance. Together, these results demonstrate that visual reconstruction from V1 wide-field calcium signals is strongly stimulus-dependent and category-dependent. Scenes featuring prominent spatial organization, stable targets, or simple background structures are inherently easier to decode, whereas complex backgrounds, high dynamic range, and subtle feature boundaries elevate reconstruction difficulty. Ultimately, these metrics indicate that the visual information accessible for mesoscopic V1 decoding varies systematically across natural scene statistics.

### 2.4 Representation alignment between visual and optogenetic signals

STAR employs a dual-branch architecture to process visual-evoked and optogenetic-evoked signals. To evaluate the specific contribution of each branch and input modality, we performed scene reconstructions independently for each signal type. Specifically, we investigated whether optogenetic-evoked V1 signals can mimic the neural representations naturally induced by visual photostimulation. Figure 5 illustrates the reconstructions generated from each modality across identical natural visual stimuli. Reconstructions derived from optogenetic-evoked signals displayed clear structural similarities to those obtained from visual-evoked wide-field signals. For stimuli featuring animal subjects, natural backgrounds, or distinct spatial layouts, both modalities successfully recovered comparable object locations and primary contours. These demonstrate that optogenetically driven cortical responses preserve meaningful spatial representation information corresponding to visual inputs.

**FIGURE 5.**
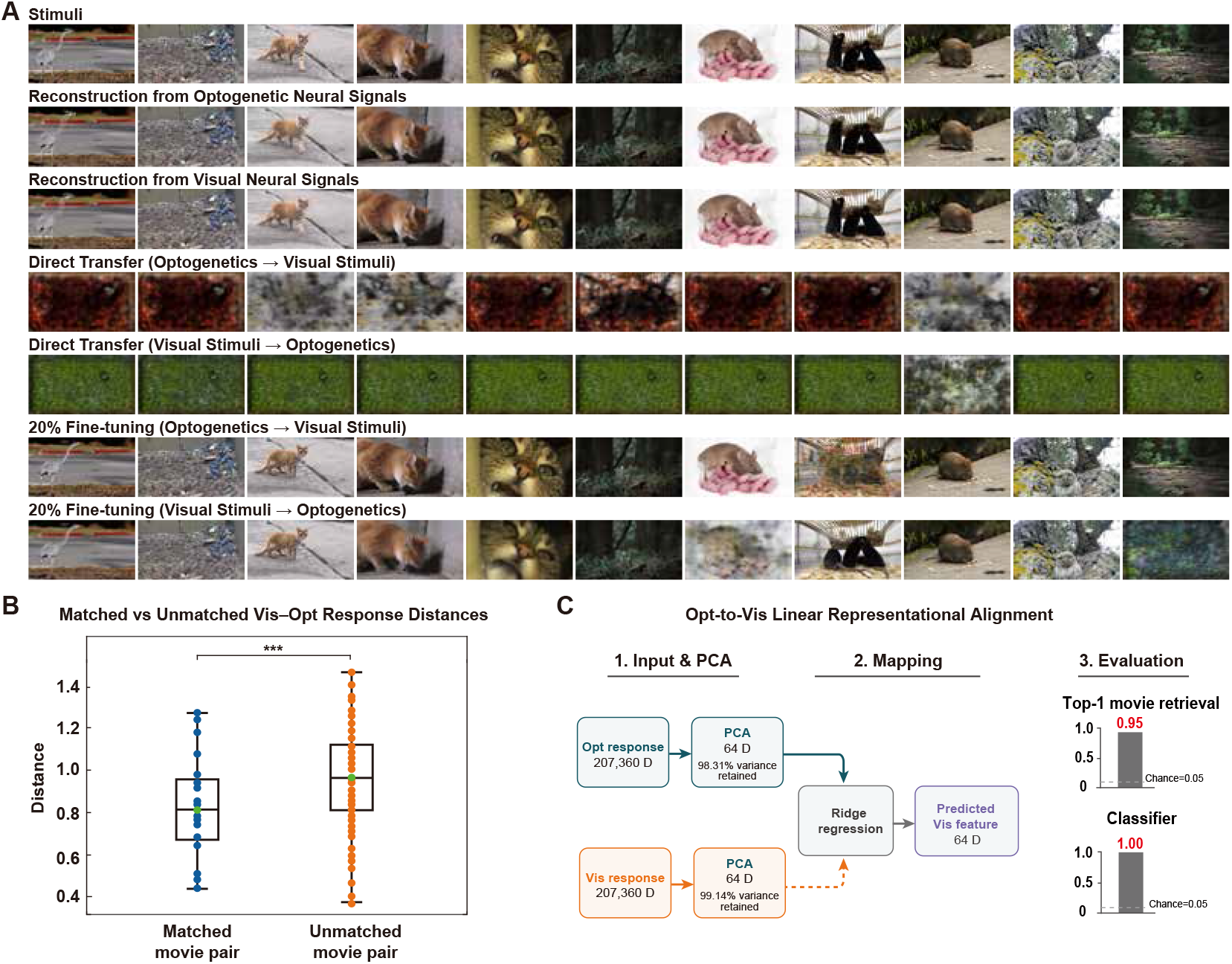
Cross-modality reconstruction between visual-evoked and optogenetic-evoked V1 signals. (A) Single-modality models reconstructed natural scenes from both signal types, whereas direct cross-modality transfer produced degraded images. Reconstruction quality was largely recovered after fine-tuning with 20% target-modality data. (B) Matched visual–optogenetic response pairs showed smaller distances than unmatched pairs. (C) Linear mapping from optogenetic to visual response features enabled accurate movie retrieval and classification, indicating partial alignment between the two response spaces.

Despite these similarities, the two response modalities are not fully equivalent (Figure S4). Direct cross-domain transfer of single-modality models led to severe performance degradation (Figure 5A). Both optogenetic-to-visual and visual-to-optogenetic direct transfers resulted in blurred reconstructions with compromised structural integrity. However, fine-tuning the model with a small fraction (20%) of target-domain data substantially restored reconstruction quality, making primary visual features recognizable once again (Figure 5A). These results indicate that visual-evoked and optogenetic-evoked V1 signals largely share foundational visual representations.

To further quantify this representation overlap, we evaluated response distances between matched and unmatched movie pairs across modalities (Figure 5B). Distance metrics for matched movie pairs were significantly lower than those for unmatched pairs, confirming that visual and optogenetic responses driven by identical stimulus content share higher representational similarity. This finding demonstrates a content-specific representational alignment at the response-distance level. Furthermore, we tested the linear alignability of the two response spaces (Figure 5C). We dimensionality-reduced responses from both conditions via Principal Component Analysis (PCA) and applied ridge regression to map optogenetic-evoked features into the visual-evoked space. The transformed representations were evaluated using movie retrieval and classification tasks, performing well above chance level. These indicate that although optogenetic stimulation does not perfectly replicate visually evoked neural patterns, the domain discrepancy can be partially reconciled via a linear transformation. Together, these findings demonstrate that visual-evoked and optogenetic-evoked V1 signals share partially aligned representational spaces. Optogenetic pattern stimulation effectively captures core features of natural visual inputs while retaining distinct distribution characteristics, providing the rationale for deploying a dual-branch network to achieve unified reconstruction.

## 3 Discussion

In this study, we developed a novel dual-modal framework in mouse V1, demonstrating that arbitrary natural movie frames can be translated into spatiotemporally patterned optogenetic inputs that evoke highly comparable, content-related cortical states. Our neural decoding results provide critical insights into the visual representations embedded within wide-field macroscale dynamics. We demonstrated that the visual-evoked mesoscopic signals from V1 contain rich structural, layout, and luminance properties that enable successful pixel-level and structural reconstruction of dynamic natural scenes. Furthermore, STAR demonstrated stable reconstruction fidelity even when operating on single-trial recordings, confirming that the decoding reliance is highly specific to visual stimulus-related neural activity localized in V1. Crucially, our work bridges a crucial domain gap between endogenous photostimulation and artificial network driving. We demonstrated that visual-evoked and optogenetic-evoked V1 signals share a partially aligned representational geometry, sharing spatial contours and content-related structures that allow accurate movie retrieval and classification via simple linear mapping. To reconcile the distinct distribution differences and cross-modality degradation discovered during direct model transfers, our unified dual-branch STAR framework successfully utilized separate input encoders to capture modality-specific feature discrepancies while mapping them into a shared reconstruction space. By confirming that complex natural visual information can be both naturally processed and synthetically injected into V1 circuits with high decoding fidelity, this work validates the use of high-dimensional patterned optogenetics for sensory emulation. Ultimately, this paradigm establishes a robust computational and experimental architecture for closed-loop, bidirectional brain-machine interfaces and broadens the horizons for investigating large-scale neocortical processing.

### 3.1 Interpretation of neuronal response to natural movies

A persistent frontier in sensory neuroscience is expanding foundational coding principles derived from simple, low-dimensional artificial stimuli to complex, ethologically relevant natural scene statistics (29; 17). Historically, the functional properties of V1 have been categorized using low-dimensional artificial stimuli, such as drifting sinusoidal gratings or localized sparse noise, which emphasize classical receptive field (RF) characteristics like orientation selectivity and spatial frequency tuning (30). However, natural scenes are governed by highly complex spatiotemporal correlations, scale invariance, and nonlinear contextual modulations that classical linear-nonlinear models fail to capture fully (29; 31), leaving a critical knowledge gap regarding how those hardwired features orchestrate distributed representations during natural vision. By establishing that wide-field macroscale dynamics in mouse V1 preserve enough structural, layout, and global luminance information to decode dynamic natural video frames, our study demonstrates a close alignment between laboratory-mapped RFs and complex scene statistics. Crucially, our ability to translate arbitrary natural movie frames into spatiotemporally patterned optogenetic arrays, and subsequently decode them under a unified model framework (STAR), proves that this artificial-to-natural gap can be bridged not only observationally but also engineered causally via precise optogenetic pattern injection.

The success of our STAR in reconstructing natural video frames directly from V1 wide-field signals reveals that mesoscopic neocortical activity retains a high degree of structural, layout, and global luminance information regarding the ongoing visual scene. This finding supports the principle of efficient population coding, which posits that the visual system organizes sensory representations to compress redundancy while preserving critical components of environmental features (32). Interestingly, our observation that reconstruction accuracy exhibits clear stimulus and category dependence, where scenes with clear object boundaries or simple backgrounds (e.g., mouse and natural scene categories) yield higher structural similarity (SSIM) and pixel fidelity (PSNR), suggesting that V1 networks prioritize stable spatial contours and macroscopic layouts over high-frequency local details during natural vision (33; 34). This is further corroborated by our feature analysis, which demonstrated that no single low-level visual attribute (such as motion energy, edge density, or RMS contrast) completely accounts for decoding variance, indicating that V1 channels integrate joint spatiotemporal features nonlinearly.

Furthermore, our control experiments contrasting V1 and non-V1 regions confirm that this natural scene representation is heavily anchored to retinotopically defined sensory coordinates. While wide-field calcium imaging aggregates mesoscopic signals across thousands of neurons, including local neuropil signals, axonal projections, and diverse cellular layers, the distinct performance degradation observed when using non-V1 signals emphasizes that macroscale cortical dynamics are not merely dominated by non-specific global arousal or widespread behavioral movements (24; 35). Instead, structured sensory information is precisely localized within V1 networks. The stable decoding performance preserved under single-trial conditions, despite trial-to-trial fluctuations, underscores that the population code for natural scenes is highly distributed and redundant, ensuring reliable information transmission even during ongoing endogenous brain-state fluctuations.

Crucially, our interpretation of natural movie responses extends beyond passive observation to active engineering via optogenetic pattern injection. The observation that spatiotemporally converted optogenetic movie stimulation evokes V1 states with representational geometry tightly aligned with actual visual photostimulation confirms that the underlying cortical architecture handles artificial spatiotemporal inputs through a matching low-dimensional manifold (36). While a significant domain gap manifests when directly transferring decoding models between the two modalities due to different spatial distributions and target opsin-expression boundaries, the success of a linear transformation (via ridge regression) in achieving high movie retrieval and classification above chance level proves that the synthetic representations remain fundamentally content-related. By demonstrating that synthetic patterns can be injected to replicate the macroscopic temporal and spatial dynamics of natural scene processing, this study indicates that V1 operates under hardwired spatiotemporal constraints that shape both endogenous and artificially driven neural spaces.

### 3.2 Functional significance and the underlying neural mechanism

We find that spatiotemporally converted optogenetic patterns can evoke mesoscopic cortical patterns highly comparable to endogenous visual responses. This demonstrates that V1 networks possess hardwired physiological boundaries, or low-dimensional manifolds, that constrain population activity (36). Rather than acting as an unconstrained, infinite-dimensional processing substrate, the local microcircuitry of V1 routes incoming sensory arrays through stereotypical spatiotemporal pathways. When artificial, pattern-translated optogenetic currents are injected directly into the network via the ultra-sensitive opsin rsChRmine, they successfully align with these pre-configured manifolds. This explains why simple linear transformations, such as ridge regression, achieve accurate cross-modal movie retrieval and classification significantly above chance levels, confirming that the underlying neural representation remains fundamentally content-bound regardless of the upstream stimulus source.

However, the distinct domain gap revealed by our cross-modality model transfer experiments, wherein directly testing a visual-trained model on optogenetic responses (and *vice versa*) causes substantial reconstruction degradation, points to critical mechanistic differences between sensory photostimulation and direct optogenetic driving. Photostimulation relies on the classic, hierarchical retinogeniculate pathway, entering V1 primarily through thalamocortical inputs terminating in Layer 4 before propagating through complex local feedforward and feedback canonical circuits (37). This biological routing inherently includes extensive synaptic filtering, recurrent network normalization, and recruitment of local inhibitory microcircuits (e.g., parvalbumin- and somatostatin-positive interneurons) that sculpt the temporal tuning and spatial boundaries of the response (38). In contrast, our pattern-converted optogenetic stimulation bypasses the pre-cortical visual pathway and directly depolarizes the membranes of the target V1 excitatory neurons expressing rsChRmine. This method bypasses natural synaptic delays, alters standard spatial receptive field summation boundaries, and can lead to a less refined spatial distribution due to target expression boundaries and light scatter within the cortical tissue (**?** 25).

The success of our specialized dual-branch STAR architecture provides a machine-learning proof of concept for how the brain might naturally reconcile these modality-specific distribution shifts. By assigning individual, isolated input branches to extract specific features from each stimulation condition and mapping them into a unified, shared decoding space, the model overcomes individual domain discrepancies without sacrificing the core visual content. In a physiological context, this shared decoding space resembles how downstream higher visual areas or polymodal motor cortices must extract robust semantic meaning from V1 arrays despite ongoing internal state variations, shifts in global arousal, or unexpected feedforward input variations (24; 35).

Furthermore, our comprehensive ablation analysis underscores the importance of treating cortical dynamics as integrated spatiotemporal structures rather than isolated frames. Removing the TRCNN or replacing it with uncoupled 1D or 2D convolutions severely degraded reconstruction metrics. This decline highlights that the functional readout of V1 activity relies heavily on the temporal integration of successive spatial states. The complementary contributions of our spatial and temporal attention mechanisms mirror the known physiological dependencies of V1 on local recurrent amplification and long-range temporal smoothing, which help maintain visual persistence and contextual continuity during rapid natural viewing (39). Ultimately, our results suggest that the functional significance of V1 macroscale dynamics during natural vision lies in its capacity to form low-dimensional, spatiotemporally robust manifolds. These manifolds compress complex environmental features into stable population codes that are highly resistant to input anomalies and easily read by downstream networks.

### 3.3 Implications for visual cortical prostheses and future work

This successful translation from natural scenes into patterned optogenetic currents carries major implications for the design of next-generation visual neuroprostheses and visual restoration strategies. For decades, traditional visual restoration clinical trials have relied primarily on microelectrode arrays implanted in the retina or directly into the primary visual cortex (40; 41). While these electrical prostheses have succeeded in restoring basic phosphene-based visual sensations, they suffer from fundamental biophysical constraints: electrical current spreads non-selectively through the parenchymal extracellular matrix, leading to large, overlapping artificial phosphenes, a lack of cell-type specificity, and tissue damage over time due to high charge-density delivery (42). By utilizing targeted optogenetic actuators, such as the red-shifted, ultra-sensitive microbial channelrhodopsin rsChRmine (26; 27), our paradigm highlights the superiority of optical manipulation over electrical stimulation. Optogenetics offers exquisite cellular selectivity, preventing the non-specific activation of passing axons and allowing for much higher spatial resolution corresponding directly to pixels or features within a structural natural frame (25).

Evaluating our approach alongside other emerging optogenetic therapies highlights the unique strengths of this paradigm. Early clinical milestones in optogenetic visual restoration have targeted the retina, delivering channelrhodopsin vectors to retinal ganglion cells via intravitreal injections to treat severe outer-retinal degeneration like retinitis pigmentosa (43). However, these retinal interventions remain entirely dependent on an intact optic nerve and downstream subcortical circuitry. For patients suffering from profound optic nerve trauma, advanced glaucoma, or bilateral cortical blindness, restoring vision requires directly addressing the neocortex (40). Our paradigm addresses this need by demonstrating that complex, dynamic sensory inputs can be successfully injected directly into V1, bypassing pre-cortical pathways entirely while generating high-fidelity representations that downstream networks can read.

Furthermore, the design of our STAR offers a valuable computational blueprint for optimizing real-time visual prostheses. Any device meant to restore vision must translate camera-acquired visual scenes into structured stimulation commands. However, as our cross-modality model transfer experiments reveal, there is a clear domain gap between biological sensory pathways and direct cortical stimulation due to altered spatial distribution and light scatter in tissue (44). A successful cortical prosthesis cannot simply act as a naive camera-to-microscopic display array; instead, it must employ smart encoder models that adaptively pre-warp and calibrate spatial patterns to match the underlying, low-dimensional representational manifolds of the host cortex. By demonstrating that separate feature branches can map disparate stimulation domains into a unified, shared reconstruction space, this work establishes a clear path toward closed-loop, intelligent biomimetic encoders capable of delivering high-fidelity, high-dimensional visual restoration. More broadly, our work demonstrates that neural decoding provides a powerful framework for evaluating artificial sensory representations. By linking stimulation patterns to the information encoded in cortical population activity, this approach offers a principled strategy for developing and benchmarking future visual cortical prostheses while providing new opportunities to investigate how natural and artificial sensory signals are represented across large-scale cortical networks.

## 4 Materials and methods

### 4.1 Animal

Four male and six female GP4.3 mice (C57BL/6J-Tg(Thy1-GCaMP6s)GP4.3Dkim/J, JAX: 024275)

(28) aged 2–4 months were used. All experimental procedures were conducted under animal welfare guidelines and approved by the Institutional Animal Care and Use Committee at the Chinese Institute for Brain Research, Beijing. All relevant ethical regulations for animal use were followed.

### 4.2 Virus injection and cranial window implantation

To achieve widespread and even expression in the primary visual cortex (V1), adeno-associated viruses (AAVs) expressing non-floxed rsChRmine (AAV2/9-CaMKIIa-rsChRmine-oScarlet-KV 2.1-WPRE, 5.5× 10^12^ GC/ml in 1 ×PBS) were injected into the transverse sinus of GP4.3 mouse pups at postnatal day 0–1 (P0–P1) (45). Pups were removed from their home cages, anesthetized by hypothermia on ice for 2–3 minutes, and head-fixed on an ice-cooled metal plate. A small incision was then made on the left transverse sinuses, approximately 1 mm anterior and 0.5 mm lateral to the lambda, and 1000 nl of virus was injected at a rate of 20 nl/s using a glass pipette. After injection, the incisions were sealed with tissue adhesive (3M Vetbond), and the pups were allowed to recover on a warming pad for 5 minutes before being returned to their home cage.

Eight weeks after viral injection, a cranial imaging window was implanted over V1. Following attachment of a head plate to the skull, a 3-mm craniotomy centered 2.5 mm lateral and 0.5 mm anterior to lambda was performed to expose V1. A stack of three coverslips (4 mm, 3 mm, 3 mm) bonded with optical adhesive (Norland) was then placed over the craniotomy and the surrounding skull and secured with dental cement (C&B Metabond). Mice were allowed to recover for at least 7 days before imaging experiments.

### 4.3 Dual-modal optical system

To enable simultaneous wide-field calcium imaging and patterned optogenetic stimulation, we developed a custom dual-modal optical system integrating time-division multiplexed fluorescence imaging with DMD-based photostimulation (Figure 1A). For wide-field imaging, two LEDs (470 nm and 405 nm) served as excitation sources for calcium-dependent and reference fluorescence imaging, respectively. The emitted light was filtered through band-pass excitation filters (475±12.5 nm and 405±12.5 nm) and collimated using aspherical lenses. The two excitation paths were combined by a dichroic mirror (DM1: LP455) and relayed to the sample through two additional dichroic mirrors (DM2: LP495; DM3: BP 500–550 nm band-pass dichroic) and an objective lens (75 mm f/2.8).

Emitted fluorescence was collected by the objective lens, transmitted through DM3, filtered by an emission band-pass filter (520 ± 18 nm, Edmund), and imaged onto an sCMOS camera (Tucsen, Dhyana 410D) through a tube lens (TL1: f = 100 mm). Calcium-dependent (470 nm) and reference (405 nm) signals were acquired alternately using time-division multiplexing (TDM). Images were recorded at a spatial resolution of 512 ×512 pixels over a 5 mm×5 mm field of view (FOV), corresponding to 9.8 *µ*m per pixel. Each channel was recorded at 10 Hz using an exposure time of approximately 10 ms.

For patterned optical stimulation, a digital micromirror device (DMD; Fldiscovery, F6500) was used to spatially modulate a collimated laser beam (590 nm, QAXK-LASER-BYR1). The generated stimulation patterns were projected onto the same imaging plane through a second tube lens (TL2, f = 125 mm), DM2, DM3, and the objective lens. Pattern size and position were calibrated to match predefined 3 mm× 3 mm regions of interest (ROIs) within the imaging field. The projected stimulation patterns had a spatial resolution of 662× 662 pixels and a refresh rate of 60 Hz. The intensity of each projected pixel could be independently controlled over an 8-bit range (0–255). Hardware control, synchronization, and TDM image acquisition were implemented using custom software written in LabVIEW (National Instruments).

### 4.4 Visual stimulation

A 21.5-inch LED monitor was placed 15 cm from the mouse’s right eye, centered at 45° azimuth and 25° elevation, covering a visual field of 116° *×* 84°. During the habituation period, the screen displayed a gray background. To measure retinotopic maps, mice were presented with spherically corrected drifting bars containing a flickering black-and-white checkerboard pattern that drifted in the four cardinal directions (46).

To elicit robust and diverse responses in V1, a library of 20 natural movie clips (1280 × 720 pixels) was presented. Each clip lasted 5 s and was displayed at 30 Hz. The stimulus set was selected to encompass a broad range of ecologically meaningful scenarios for mice, including predator-related scenes, prey and foraging behaviors, conspecific social interactions, and natural landscapes. A complete list of movie clips and their classifications is provided in Table S1. During imaging sessions, movie clips were presented in a pseudorandom sequence, with each clip repeated three times. A uniform gray screen was displayed during a 3–5 s inter-stimulus interval.

### 4.5 *In vivo* wide-field calcium imaging data acquisition and analysis

Following recovery, mice were head-fixed and allowed to move freely on a rotating treadmill during imaging experiments. Wide-field calcium imaging and patterned optogenetic stimulation were performed simultaneously using the dual-modal optical system described above.

Hemodynamic artifacts were removed from wide-field calcium signals (47). For each channel, baseline fluorescence was estimated as the mean fluorescence during inter-stimulus intervals, and the corresponding Δ*F/F* values were calculated for the 470 nm and 405 nm channels. The 405 nm isosbestic component was then subtracted from the 470 nm signal to obtain the hemodynamic-corrected fluorescence response. To correct for slow baseline drift, a sliding-window percentile-based baseline estimation method was applied (window = 15 s, 25th percentile), and the estimated baseline was subtracted from the fluorescence traces. Finally, the traces were converted to *z* scores using the mean and standard deviation calculated from the inter-stimulus interval.

Individual retinotopic maps were obtained by analyzing neural responses to drifting bars, as described previously (48; 49). These maps were used to identify V1 and to establish the correspondence between visual and cortical coordinates for subsequent analyses and optogenetic stimulation.

### 4.6 Optogenetic stimulation

Optogenetic stimulation patterns were generated from natural movie clips using a two-stage encoding framework. First, feature encoding extracted specific visual features from the input movies. Second, retinotopic transformation projected the encoded features onto their corresponding cortical locations in V1 based on the measured retinotopic map of each mouse.

#### 4.6.1 Feature encoding

We evaluated three feature encoding strategies that capture distinct visual features: motion, object boundaries, and visual saliency. Before feature extraction, all movie frames were spatially smoothed to match mouse visual acuity.

1. **Motion**. Each frame was denoised using bilateral filtering (*σ*_*color*_ = 200, *σ*_*space*_ ∈ [25, 75, 150] depending on the video) and Gaussian filtering (*σ*_*Gaussian*_ = 5). Motion was estimated by computing temporal difference images between consecutive frames. The resulting difference images were converted to grayscale, smoothed with an additional Gaussian filter (*σ* = 5), and thresholded to remove values below 10.
2. **Object boundaries**. Each frame was Gaussian filtered (*σ* = 11) before horizontal and vertical Sobel gradients were computed. The two gradient components were combined to generate boundary maps, which were subsequently smoothed with a Gaussian filter (*σ* = 3). Boundary intensities were normalized to the range of 0–255, and values below 50 were discarded.
3. **Visual saliency**. Dynamic saliency maps were generated using the method described previously (50). Saliency values below 20 were discarded.

To evaluate the fidelity of each encoding strategy, optogenetic-evoked cortical activity was compared with visual-evoked activity using three complementary metrics: cosine similarity, Pearson’s correlation coefficient, and a soft F1 score.

##### Cosine similarity

Global similarity between visually and optogenetically evoked activity was quantified as:

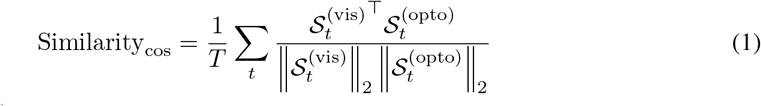

where 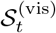 and 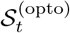 denote the vectorized cortical activity maps evoked by visual and optogenetic stimulation, respectively, at time *t*, and *T* is the total number of frames.

##### Pearson’s correlation coefficient

Pearson’s correlation coefficient was computed between visually and optogenetically evoked activity maps at each frame and then averaged across time:

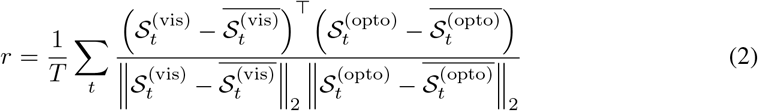

##### Soft F1 score

To quantify the spatial overlap between visually and optogenetically evoked activity, the soft intersection at each frame was defined as

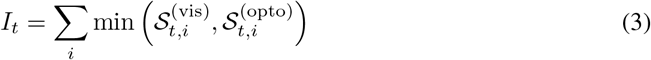

where *S*_*t,i*_ denotes the rectified *z*-score activity at cortical location *i* and time *t*. The soft precision and recall were then computed as

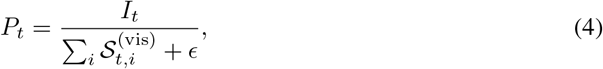

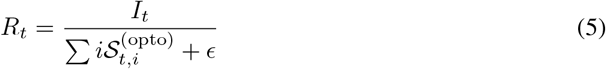

where *ϵ* is a small constant introduced to avoid division by zero. The soft F1 score was calculated as the temporal average of the harmonic mean of precision and recall:

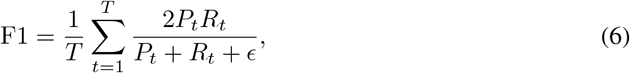

The soft F1 score ranges from 0 to 1, with higher values indicating greater similarity between visually and optogenetically evoked cortical activity patterns.

#### 4.6.2 Retinotopic transformation

The relationship between laser intensity and cortical activity was established by measuring calcium responses across a range of stimulation intensities. Cubic B-spline interpolation was used to generate a continuous intensity–response function, enabling encoded feature values to be converted into optogenetic stimulation intensities. Based on this calibration, stimulation intensities were restricted to the range of 50–225 in subsequent experiments to ensure reliable and robust cortical activation. Finally, the encoded feature maps were transformed according to each mouse’s retinotopic map to generate the final optogenetic stimulation patterns, which were projected onto the left V1.

### 4.7 STAR model for visual reconstruction

#### 4.7.1 STAR model structure

The input neural signal is denoted as *R* ∈ℝ^*T×H×W*^, where *T* is the temporal window length, and *H* and *W* represent the spatial dimensions of cortical activity. We first localize the primary visual cortex (V1) based on retinotopic mapping and crop the corresponding region as input. This step preserves relevant visual structure while reducing irrelevant noise. Given the neural signal *R* and its corresponding video frame *S*, the model learns a mapping function *f* as *Ŝ* = *f* (*R*), where *Ŝ* is the reconstructed frame. The overall STAR pipeline is

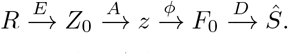

Here, *E* denotes the spatio-temporal encoder, *A* the attention aggregation module, *ϕ* the projection module, and *D* the reconstruction decoder.

1. **Spatiotemporal encoding** The encoder *E*(*·*) extracts joint spatiotemporal features from calcium signals. The input is first arranged as a 3D spatiotemporal volume. A 3D convolution layer is used to extract initial features, followed by several 3D residual blocks for deeper modeling. Each residual block uses spatiotemporal kernels. The temporal dimension can adopt different dilation rates, which enlarge the temporal receptive field and adapt to different dynamics across modalities. The encoding process is written as *Z*_0_ = *E*(*R*), where *Z*_0_ denotes the encoded features. In practice, visual-evoked and optogenetic-evoked signals are processed by two encoders, *E*^(*v*)^(*·*) and *E*^(*o*)^( *·*). They share the same architecture and hyperparameters, but their parameters are learned independently. This avoids structural bias and lets modality differences arise from data and learned representations.
2. **Spatiotemporal attention** After obtaining *Z*_0_, a spatiotemporal attention module is introduced to model the importance of different spatial locations and time steps. This allows the model to focus on neural patterns that are more relevant for reconstruction. The module consists of spatial attention and temporal attention. We first compute a spatial attention map. Channel-wise average pooling and max pooling are applied, concatenated, and passed through a lightweight 3D convolution:

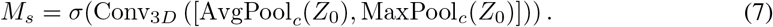

The weighted feature is *Z*_*s*_ = *Z*_0_ ⊙ *M*_*s*_. Next, we perform spatial average pooling to obtain a temporal sequence:

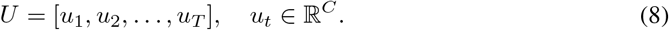

A lightweight scoring function *g*(*·*) assigns an importance score to each time step as *e*_*t*_ = *g*(*u*_*t*_), where *g*(*·*) maps each feature vector to a scalar. The attention weights are computed by softmax 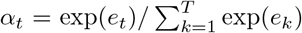. The aggregated representation is 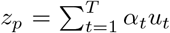. We then apply projection and normalization *z* = Norm(*P* (*z*_*p*_)). The projection head produces a 512-dimensional latent representation, which is L2-normalized before reconstruction and cross-modal alignment.

(3) **Visual reconstruction decoder** To map the latent representation to spatial features, we introduce a projection module *ϕ*(*·*). A fully connected layer reshapes the latent vector into a 2D feature map. This establishes a mapping from neural activity to visual spatial structure. We then use a shared U-Net style decoder *D*(*·*) to refine the features. The decoder takes the projected feature as input. It uses an encoder–decoder structure with skip connections. The architecture includes an initial convolution layer, two downsampling blocks, a bottleneck residual module, and two upsampling blocks. Skip connections fuse multi-scale features. This helps recover spatial structure while preserving details at different scales. The reconstruction is *F*_0_ = *ϕ*(*z*), then *Ŝ* = *D*(*F*_0_). The output is a three-channel RGB frame, constrained to [0, 1] by a sigmoid function. The decoder is shared across both modalities, so they are reconstructed in the same visual space.
(4) **Reconstruction loss** For each stimulation modality, scene reconstruction is supervised using an *L*_2_ pixel-wise mean squared error loss as *L*_rec_ = *MSE*(*Ŝ*^*m*^, *S*), *m*∈ {*v, o* }, where *Ŝ*^*m*^ denotes the reconstructed image frame and *S* represents the corresponding ground-truth visual stimulus. This objective directly penalizes spatial reconstruction errors to enforce pixel-level fidelity.

#### 4.7.2 Model training and testing

To analyze the relationship between different neural signals, a series of training and testing was conducted, including single-modality reconstruction, cross-modality transfer, incremental fine-tuning, and joint learning. Each one targets a different aspect, such as decodability, representation consistency, adaptability, and alignment.

##### Single-modality Reconstruction

To train separate models on visual-evoked and optogenetic-evoked activity, each model is trained on its own training set and evaluated on the same test set.

##### Cross-modality Transfer

A model trained on one modality is directly applied to the other, without updating parameters. Let *θ*_*s*_ be the model trained on the source modality. For a target input *R*^(*t*)^, the reconstruction is 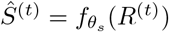. If representations are aligned, performance should remain stable. Otherwise, performance will drop, indicating a mismatch.

##### Incremental Fine-tuning

Fine-tuning of the model with a small subset of target data was done for both modalities. Let *f*_*θ*_ be the pretrained model and 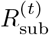 the subset. Fine-tuning gives 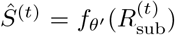. If performance recovers with few samples, it suggests a shared but misaligned representation space.

##### Multi-modal Joint Learning

Dual-branch training includes both modality losses, visual-evoked 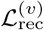 and optogenetic-evoked 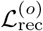 jointly, where each modality has its own encoder, while both modalities share the same reconstruction decode r. The total loss is 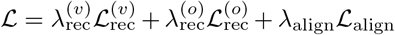. The alignment loss is 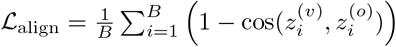. Samples are paired by the same stimulus, so alignment is applied sample-wise. In all experiments, we set 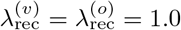 and *λ* _align_ = 0.01.

#### 4.7.3 Evaluation metrics

Reconstruction quality was evaluated using peak signal-to-noise ratio (PSNR) and the structural similarity index measure (SSIM). Both metrics were computed between each reconstructed RGB frame and its corresponding ground-truth frame. Image intensities were normalized to the range [0, 1] before evaluation. PSNR and SSIM were first calculated for individual test frames and then averaged over all test frames for each experimental condition. Higher PSNR and SSIM values indicate better pixel-level reconstruction fidelity and structural similarity, respectively.

For cross-modal stimulus retrieval, each optogenetic representation mapped into the visual representation space was compared with the candidate visual representations. Retrieval was considered correct when the best-matching candidate had the same stimulus identity as the query. Performance was reported as top-1 retrieval accuracy.

#### 4.7.4 Implementation Details

Training data consisted of temporally aligned samples paired to identical visual stimuli, with both modalities sharing the exact same target ground-truth frame. Optimization was performed using the Adam optimizer with an initial learning rate of 10^*−*4^, adjusted according to a multi-stage learning rate decay schedule. Model selection and early stopping were governed by monitoring reconstruction performance across both input branches using a combined evaluation metric. Input sequences were extracted using a temporal window length of *T* = 6. Because both stimulation modalities were recorded within the same animal, the V1 spatial patches possessed consistent spatial dimensions, eliminating the need for spatial resizing. The network was trained with a batch size of 16 for 1000 epochs. For domain-adaptation fine-tuning experiments, the model was updated using a 20% subset of target-domain samples.

## Supplemental Materials

### Sec. 1 Video stimuli and corresponding categories

**Table S1.**
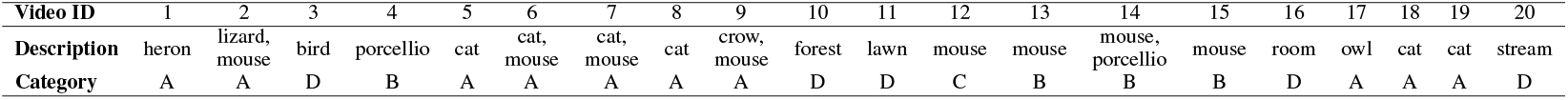
Summary of video stimuli and corresponding categories. A: predator-related scenes. B: prey and foraging behaviors. C: conspecific social interactions. D: natural landscapes.

| Video ID | 1 | 2 | 3 | 4 | 5 | 6 | 7 | 8 | 9 | 10 | 11 | 12 | 13 | 14 | 15 | 16 | 17 | 18 | 19 | 20 |
| --- | --- | --- | --- | --- | --- | --- | --- | --- | --- | --- | --- | --- | --- | --- | --- | --- | --- | --- | --- | --- |
| Description | heron | lizard, mouse | bird | porcellio | cat | cat, mouse | cat, mouse | cat | crow, mouse | forest | lawn | mouse | mouse | mouse, porcellio | mouse | room | owl | cat | cat | stream |
| Category | A | A | D | B | A | A | A | A | A | D | D | C | B | B | B | D | A | A | A | D |

**FIGURE S1.**
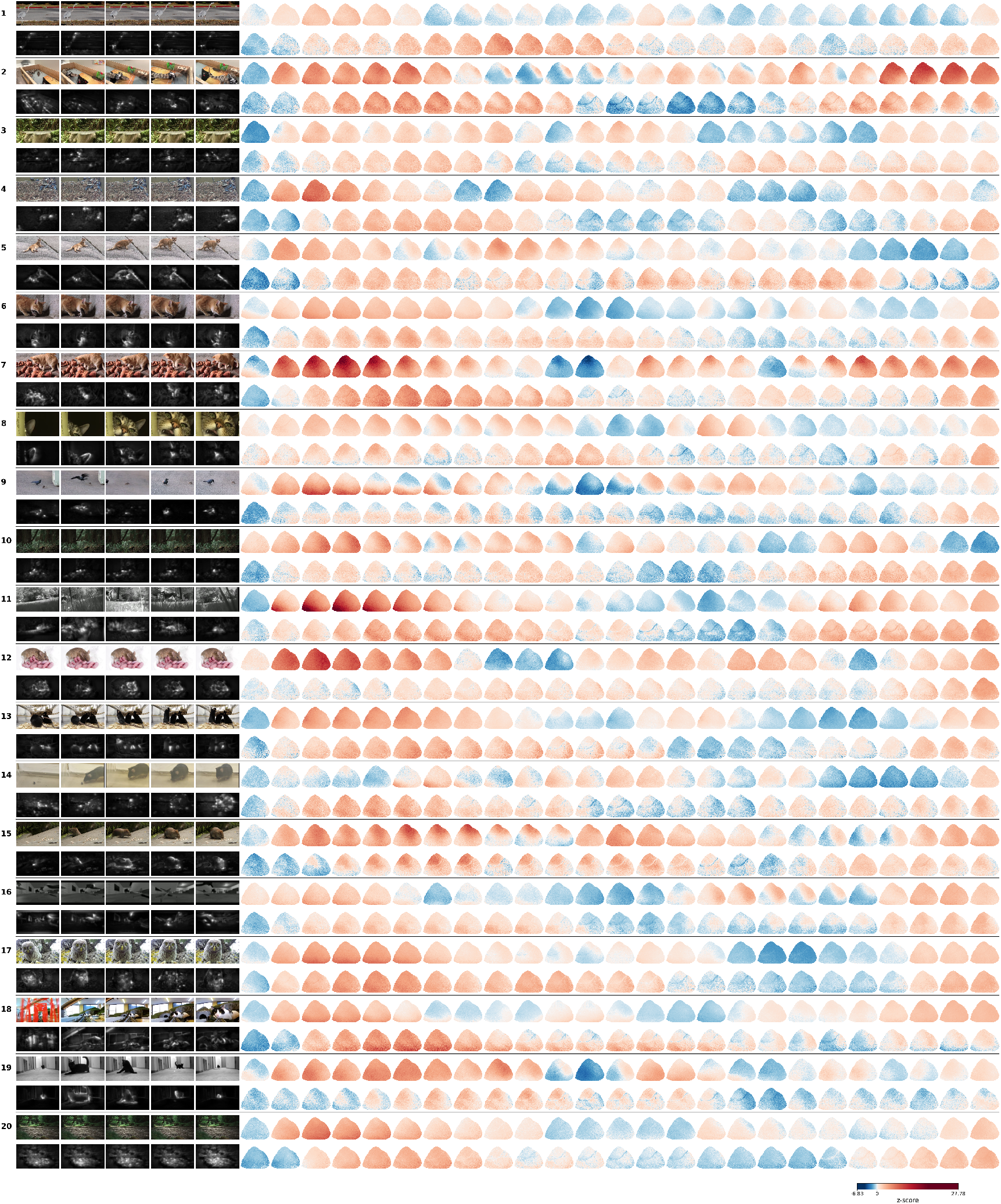
Visually and optogenetically evoked neural responses to 20 natural movies in awake mouse V1. Left, clips of nature movies (odd rows) and corresponding optogenetic stimulation patterns (even rows). Right, evoked neural responses in V1 measured using wide-field calcium imaging.

### Sec. 2 Comparing STAR with other existing models

Figure S2A shows 20 representative natural visual stimuli and their reconstructions using STAR, compared with other existing models. Overall, STAR was able to recover the main structure of the original visual scenes from V1 activity, including object locations, scene layout, and global luminance patterns. For stimuli containing animals, natural backgrounds, and complex scenes, the reconstructed images generally preserved the main regions and spatial contours of the original stimuli. In contrast, the reconstructions produced by the existing methods, SID and WISA, were overall more blurred, with less stable object structures and local boundaries. The STAR reconstructions were visually closer to the original stimuli, suggesting that V1 wide-field calcium signals contain visual information that can be used for natural scene reconstruction.

We further compared the reconstruction performance of different methods using quantitative metrics. Figure S2B shows the peak signal-to-noise ratio (PSNR) and structural similarity index measure (SSIM) results of STAR and the existing methods across different stimuli. Most data points indicate that STAR achieved higher reconstruction scores than the comparison methods, suggesting better pixel-level fidelity and structural similarity. The reconstruction accuracy still varied across stimuli, indicating that the content of natural scenes affects reconstruction difficulty. Nevertheless, STAR more consistently decoded natural visual scenes from V1 wide-field calcium signals.

**FIGURE S2.**
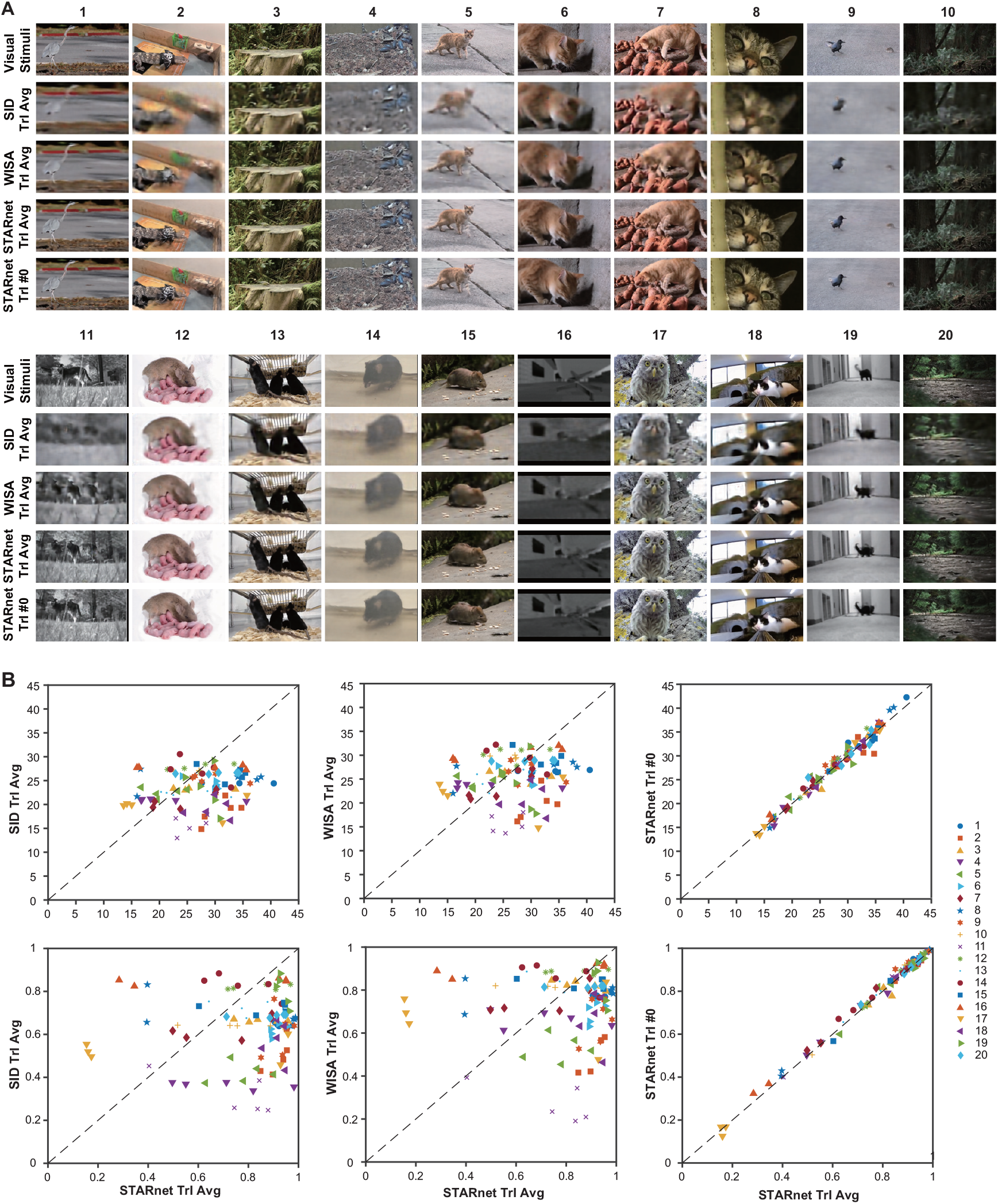
Reconstruction of natural visual scenes from visual-evoked V1 wide-field calcium signals. (A) Representative visual stimuli and reconstructions from SID, WISA, and STAR using trial-averaged inputs, together with STAR reconstructions from a single trial. (B) Quantitative comparison of reconstruction performance across stimuli using PSNR (top) and SSIM (bottom). Scatter plots compare STAR with SID and WISA under trial-averaged inputs, and compare STAR single-trial reconstruction with trial-averaged reconstruction. Dashed lines indicate equal performance.

### Sec. 3 STAR decoding with single-trial neural activity

We compared single-trial reconstruction with trial-averaged reconstruction. Trial-averaged calcium signals reduce noise from individual recordings and therefore usually provide a more stable input for the model. In contrast, single-trial signals contain greater trial-to-trial variability, making reconstruction more challenging. Table S2 reports the quantitative results of single-trial reconstruction. The relatively small variation across trials suggests that STAR does not rely solely on trial-averaged responses, but can also maintain stable reconstruction performance from single-trial V1 activity.

**Table S2.** STAR Single-trial Reconstruction.

| Trial # | Visual-evoked |  | Optogenetic-evoked |  |
| --- | --- | --- | --- | --- |
|  | PSNR↑ | SSIM↑ | PSNR↑ | SSIM↑ |
| 0 | 27.200 | 0.803 | 27.830 | 0.810 |
| 1 | 27.886 | 0.816 | 28.271 | 0.822 |
| 2 | 27.441 | 0.804 | 27.838 | 0.810 |

### Sec. 4 Cortical-region control experiment

To test whether the reconstruction specifically depends on visual-related signals from V1, we performed a cortical-region control experiment. Figure S3A marks the locations of the V1 and non-V1 regions, and Figure S3B compares reconstruction performance using signals from these two regions as input. The results show that reconstructions based on V1 signals clearly outperformed those based on non-V1 signals in both PSNR and SSIM, with the difference being particularly evident for SSIM. This indicates that the model does not achieve the same reconstruction quality from arbitrary cortical wide-field signals, but instead relies mainly on visual stimulus-related neural activity in V1. Taken together, these results show that STAR can reconstruct natural visual scenes from V1 wide-field calcium signals in awake mice and outperforms existing methods overall. The single-trial results indicate that the model retains stable reconstruction ability even from individual neural responses. The comparison between V1 and non-V1 regions further supports that this reconstruction depends on V1-specific visual information rather than nonspecific activity in wide-field signals.

**FIGURE S3.**
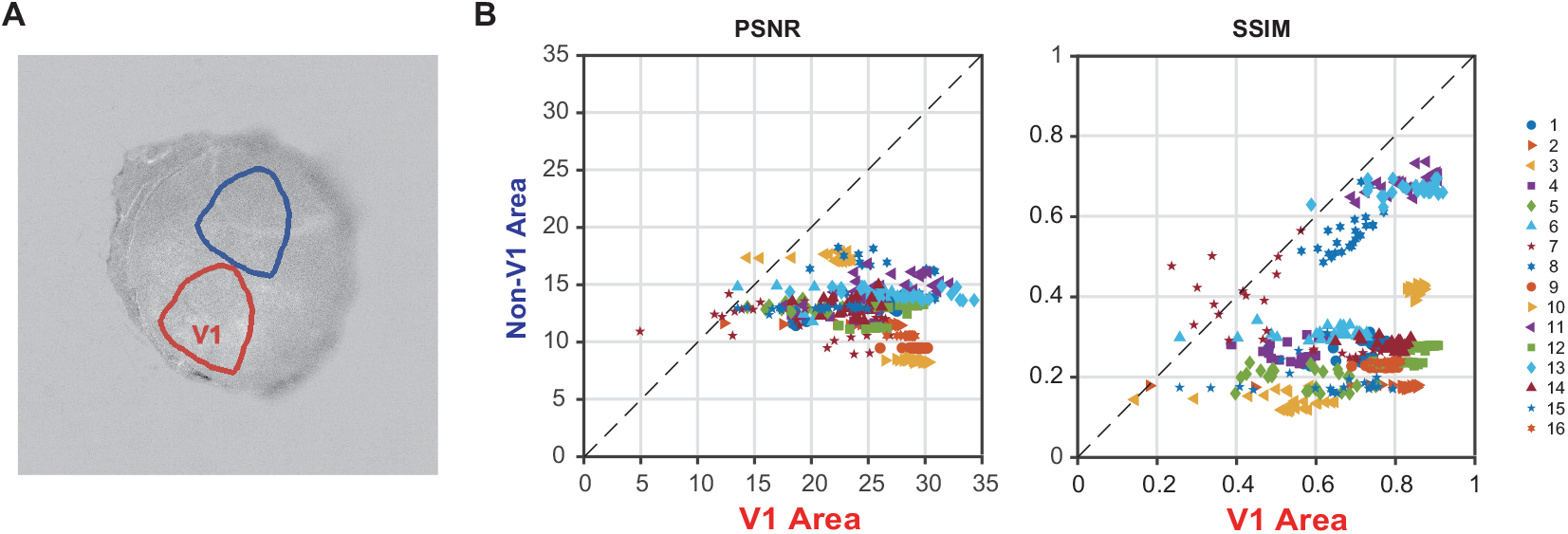
V1-specific contribution to visual reconstruction. (A) Example masks of V1 and non-V1 cortical regions. (B) Reconstruction performance using V1 signals compared with non-V1 signals, measured by PSNR and SSIM. The higher performance from V1 activity indicates that visual reconstruction relies on retinotopically defined V1 signals.

### Sec. 5 Ablation analysis identifies key components of STAR

To analyze the contribution of different designs in STAR to reconstruction performance, we conducted ablation experiments to examine the role of the main architectural components in STAR. Table S3 reports the component ablation results under the visual-evoked reconstruction setting. We also compared alternative convolutional structures for replacing TRCNN. Replacing TRCNN with 2D convolution reduced the performance, while replacing it with 1D convolution caused a more pronounced decrease. This indicates that using convolution along only the spatial or temporal dimension is insufficient to capture the joint spatiotemporal information in V1 wide-field calcium signals, whereas TRCNN is better suited for the current neural signal reconstruction task.

**Table S3.** Ablation Study on the Proposed Model Components.

| **Ablation Settings** | **PSNR↑** | **SSIM↑** |
| --- | --- | --- |
| Full Model | 27.654 | 0.814 |
| w/o TRCNN | 27.190 | 0.808 |
| w/o Spatial Attention | 27.477 | 0.810 |
| w/o Temporal Attention | 27.444 | 0.811 |
| Replace TRCNN with 2D Convolution | 27.432 | 0.809 |
| Replace TRCNN with 1D Convolution | 26.463 | 0.783 |

### Sec. 6 Movie-Wise Representational Topography in t-SNE Space

To further examine the fine-grained representational structure of visual-evoked and optogenetic-evoked V1 neural activity across individual stimuli, neural response samples from both modalities were embedded into a shared two-dimensional t-SNE space and partitioned by movie identity (Figure S4). For each of the 20 natural movies (Mov 1–Mov 20), visual-evoked responses (orange triangles) and optogenetic-evoked responses (blue filled circles) exhibit distinct clustering behavior within the shared embedding space. Visual-evoked dynamics consistently form compact, well-defined clusters that reflect the underlying temporal continuity and spatial statistics of the natural video inputs. In contrast, optogenetic-evoked responses form corresponding yet spatially shifted clusters, revealing an ongoing domain gap between direct cortical optogenetic driving and endogenous visual photostimulation. Despite this spatial separation between modalities, the relative internal geometry of the responses for each movie is preserved. Within individual panels, optogenetic-evoked samples remain localized in specific sub-regions of the manifold that mirror the trajectory of their visual-evoked counterparts rather than dispersing randomly across the t-SNE coordinate plane. This movie-wise clustering behavior demonstrates that patterned optogenetic stimulation drives V1 activity along content-specific manifolds, maintaining consistent representational topology across individual dynamic stimuli.

**FIGURE S4.**
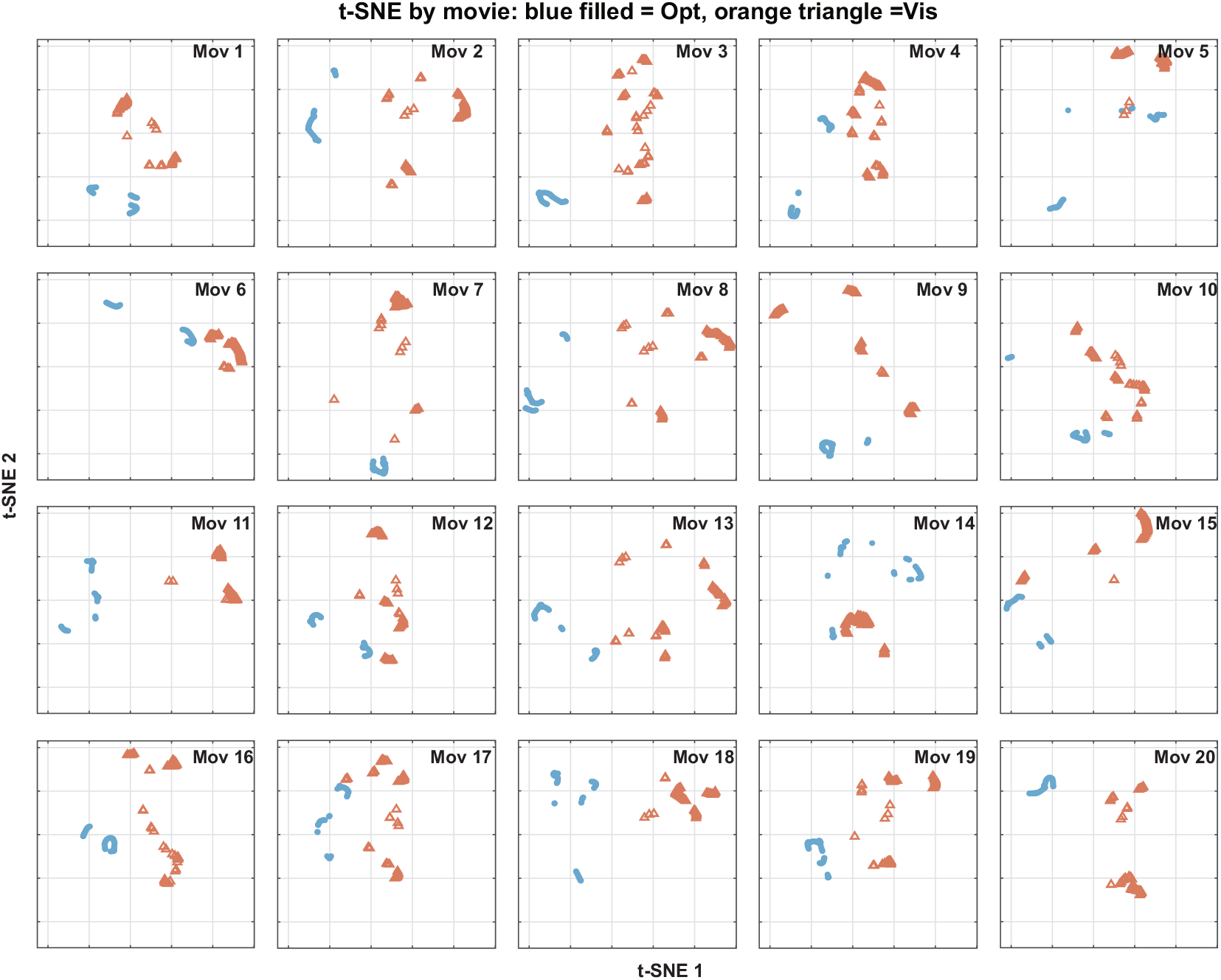
Movie-wise t-SNE visualization of visual-evoked and optogenetic-evoked V1 responses. Neural-response samples from the two modalities were embedded in a common two-dimensional t-SNE space and displayed separately for each of the 20 natural movies.

